# Association of *SRB1, ITGB2* gene polymorphisms with coronary heart disease in Chinese Han population

**DOI:** 10.1101/402792

**Authors:** Yuan Sun, Tian Long-Wang, Yong Zeng, Feng-Ying Gong, Hui-Juan Zhu, Hui Pan, Ying Wang, Jia-Li Wang

**Author notes:** Corresponding author: Tian Long-Wang.

## Abstract

**Background:** Previous studies in mice and humans have implicated the lipoprotein receptor SRB1 in association with atherosclerosis and lipid levels. In our previous proteomics research, the expression of ITGB2 has differences between epicardial and subcutaneous adipose tissue. However, the association between the reported variants and risk of coronary heart disease (CHD) was not confirmed.

**Methods:** We conducted a case–control study consisted of 496 CHD patients and 367 controls. The two groups are adjusted for age, sex, body mass index, diabetes status and the proportion of dyslipidemia. The genotypes and allele frequency of variants rs838880,rs5888,rs5889 in SRB1 and rs235326,rs2070947,rs2070946 in ITGB2 were determined using Sequenom Mass-ARRAY technology.

**Results:** The genotypes frequencies of all the six SNPs were consistent with Hardy-Weinberg Equilibrium test. For gene SRB1 rs838880, there was a significant difference in the alleles frequency(p=0.017), genotype frequency(p=0.0028), recessive model (p=0.000672) between CHD group and control group. For gene ITGB2 rs2070947, there was a significant difference in the recessive model (p=0.03). By comparing the clinical and serum metabolic indexes of SNP sites by genotype we find that among three genotypes of SRB1 rs5888,there were significant difference in the level of dyslipidemia history and serum LPA, among three genotypes of ITGB2 rs235236,there were significant difference in the levels of serum HDL,APOA1 and hypertension history, among three genotypes of ITGB2 rs2070947,there were significant difference in the level of serum APOA1,hsCRP.

**Conclusions:** Our findings indicated that SNP rs838880 of gene SRB1 and rs2070947 of gene ITGB2 are associated with the risk of CHD in Chinese han population.

## Introduction

Coronary heart disease (CHD) is the leading cause of death worldwide ^[1]^. At present a large number of clinical research and epidemiological investigations show that coronary heart disease is a complex polygenic hereditary disease, which causes include environmental factors, genetic factors and the interaction between them ^[2]^. Recent genome-wide association studies (GWAS) have identified SNPs at several loci such as chromosomes 1p13, 3q22, 6p24, 9p21 and 15q22 that are associated with risk of coronary heart disease ^[2–4]^. Scavenger receptor class B, type 1 (SRB1) is a key regulator of high density lipoprotein (HDL) metabolism. It facilitates the efflux of cholesterol from cells in peripheral tissues to HDL and mediates the selective uptake of cholesteryl esters from HDL in the liver.

SRB1 deficiency in the arterial wall results in de-regulation of vascular lipid homeostasis and induction of inflammation culminating in atherosclerosis ^[16]^. There have been reports of SNP of SRB1 genes related to atherosclerosis and coronary heart disease. But the majority of these sites were found in the European population, there was less research in Chinese population ^[5]^. Inflammation is another important mechanism of atherosclerotic and adhesion molecules plays an important role in the process of leukocyte adhesion ^[17]^. Integrin beta 2 (ITGB2), or CD18, is an important member of the integrin family in the adhesion molecule. In our early proteomics study, the expression level of ITGB2 was raised in the pericardial fat, while the expression in subcutaneous fat was significantly lower. Prove that ITGB2 may be involved in the inflammatory process of the developing of coronary heart disease ^[18]^. But there are few studies on the relationship between the single nucleotide polymorphisms and coronary heart disease in the ITGB2 gene.

## Methods

### 1. Subjects

All participants in our study were selected from the CHD research database of Peking union medical college hospital who had coronary angiography in this hospital between January 2009 and January 2015. All subjects were of self-reported Chinese Han ancestry. All subjects underwent a review of their medical history, a physical examination, and a laboratory assessment of cardiovascular risk factors.

Patients were diagnosed with CHD using coronary angiography, which revealed ≥ 50% stenosis of one or more branch of right or left main trunk, left anterior descending, left circumflex branch. We select patients with stenosis ≥75% into CHD group and coronary angiography results no abnormal or stenosis <25% into control group. The two groups are matched of gender, age, BMI, proportion of diabetes, proportion of dyslipidemia.

Hypertension diagnosed according to U.S. JNC-VI guide standard ^[19]^, two consecutive in the resting state for systolic blood pressure (SBP) ≥140 mmHg and/or diastolic blood pressure (DBP) ≥ 90 mmHg or there is a clear history of high blood pressure. Diagnose standard of diabetes mellitus is according to the American diabetes association ^[20]^, for 8h fasting venous blood sugar after meals ≥ 7.0 mmol / L and (or) 2 h venous blood sugar after meals ≥ 11.1mmol / L, or there is a clear history of type 2 diabetes. According to the WHO’s standardization of the smoking status survey method ^[21]^, a regular smoker is defined as smoking one or more cigarettes a day and smoking for six months.

### 2. SNP Selection and Genotyping

We Screening SNPs loci according to genome-wide association study (GWAS) results and literature reported. Select those has positive reported results or trends. We selected six SNPs (rs838880, rs5888, rs5889, rs235326, rs2070947, rs2070946), which have been reported but have not yet been tested in a Chinese population. SNPs with minor allele frequencies (MAF) > 5% in the HapMap database were selected. A E.Z.N.A.TMBlood DNA Midi Kit (OMEGA company) was used to extract genomic DNA from whole blood samples and Sequenom MassARRAY SNP detection platform was used to test. Detection process mainly includes primers design and synthesis, PCR amplification reaction, SAP, purification, single base extension, mass spectrometry analysis, test results using TYPER 4.0 software (sequenom).

### 3. Statistical analysis

We used SPSS 17.0 statistical packages to perform the statistical analyses and P < 0.05 was considered statistically significant. A t-est and Chi-squared test were performed to compare measuring or count index between cases and controls. Goodness.offitchi.square test was applied to each SNP in the controls to test for departure from the Hardy-Weinberg Equilibrium (HWE). Odds ratios (ORs) and 95% concidence intervals (CIs) for the allele and genotype frequencies were calculated using the Pearson Chi-square test and adjusted for gender, age, waist circumference, diabetes history, hypertension history. PLINK software 1.07 was used to assess SNP associations with CHD risk in different genetic models (dominant, recessive). Pairwise linkage disequilibrium and haplotype constructions were performed using HAPLOVIEW 4.0.

## Results

### 1. Characteristics of subjects

Our population group consisted of 496 patients with CHD and 367 controls. The CHD group including 256 man and 244 women, average age is 60.19±9.8, whereas control group including 181 man and 186 women, average age is 59.39±11.00. The CHD and control groups were age, sex, BMI, waist circumference, blood pressure, diabetes rate and dyslipidemia rate matched (p > 0.05). Baseline characteristics of the subjects are shown in Table1. The average BMI of CHD group was 26.12kg/m^2^, and 25.81kg/m^2^ in control group, both of which were slightly higher than normal crowd. In addition, the diabetes rate and dyslipidemia rate in two groups were higher than normal population, which was conform to the pathogenesis of coronary heart disease. Two groups have significant difference in family history of cardiovascular disease, 62% in CHD group and 49.2% in control group (P < 0.01), it also prompts that genetic factors plays a role in the process of the onset of coronary heart disease (CHD). The level of hsCRP in CHD group was significantly higher than that in the control group (7.4 mg/L vs 4.36 mg/L, P = 0.026), which may related to the inflammatory mechanism of coronary heart disease. In our study, the coronary heart disease group included 125 cases of stable angina pectoris (25%), 187 cases of unstable angina pectoris (37.4%), 186 cases of myocardial infarction (37.2%). Thus the acute coronary syndrome (unstable angina pectoris + myocardial infarction) accounting for 75%.

**Table 1.** Baseline Characteristics of the Subjects

|  | Controls<br>(n=367) | CHD patients<br>(n=496) | P-value |
| --- | --- | --- | --- |
| Age | 59.39±11.00 | 60.19±9.89 | 0.28 |
| Males (%) | 49.3% | 51.6% | 0.54 |
| BMI(kg/m<sup>2</sup>) | 26.12±3.79 | 25.81±3.34 | 0.54 |
| Waist (cm) | 93.58±10.28 | 91.71±14.25 | 0.37 |
| Systolic BP(mmHg) | 128.11±17.81 | 129.19±19.65 | 0.36 |
| Diastolic BP(mmHg) | 75.26±12.57 | 74.54±12.61 | 0.46 |
| Diabetes (%) | 26.7% | 30.7% | 0.188 |
| Hypertension (%) | 57.8% | 71% | <0.01 |
| Dyslipidemia (%) | 59.4% | 62% | 0.502 |
| Family history | 49.2% | 62% | <0.01 |
| Smoking/drinking (%) | 39.5% | 50.2% | <0.01 |
| Statins use (%) | 29% | 56% | 0 |
| TC (mmol/L) | 4.30±1.00 | 4.31±1.08 | 0.95 |
| TG (mmol/L) | 1.66±1.21 | 1.90±1.88 | 0.16 |
| HDL (mmol/L) | 1.12±0.43 | 1.02±0.25 | <0.01 |
| LDL (mmol/L) | 2.58±0.82 | 2.61±0.89 | 0.75 |
| APOB2 (mmol/L) | 0.92±0.12 | 0.91±0.13 | 0.66 |
| LPA (mmol/L) | 4.65±1.14 | 4.87±1.17 | 0.05 |
| cTnI (ug/L) | 0.84±7.08 | 7.86±42.51 | <0.01 |
| NT-proBNP (ug/L) | 372.32±957 | 1150.71±3675.16 | <0.01 |

apoB: apolipoprotein B, HDL: high-density lipoprotein, LDL: low-density lipoprotein, LPA: lipoprotein, TG: triglyceride, TC: total cholesterol, cTnI: cardiac troponin I $p < 0.01$ indicates statistical significance.

### 2. SNPs site Hardy-Weinberg balance test

In the controls, all SNPs were in Hardy-Weinberg equilibrium (HWE) (Table 2).

**Table 2.** Hardy-Weinberg balance of each gene site

| Gene | SNP site | chromosome | p-value |
| --- | --- | --- | --- |
| SRB1 | rs838880 | 12 | 0.68 |
|  | rs5888 | 12 | 0.45 |
|  | rs5889 | 12 | 0.39 |
| ITGB2 | rs235326 | 21 | 0.51 |
|  | rs2070947 | 21 | 0.36 |
|  | rs2070946 | 21 | 0.41 |

### 3. Case-control analysis of SNPs sites in SRB1 and ITGB2 genes

For gene SRB1 SNP rs838880, the allele frequency distributions differed between cases and controls. (*p* = 0.017, OR: 0.77, 95%CI: (0.64-0.95), After multiple logistic regression analysis corrected for gender, age, waist circumference, diabetes history, hypertension history, P = 0.01, still have significant difference. Namely C allele loci is 0.77 times of T allele loci the risk of coronary heart disease (CHD) is, C allele on coronary heart disease have protective effect. There was no significant difference in other SNP loci (Table 3). We also compared the genotype frequencies between cases and controls. For gene SRB1 SNP rs838880, the genotype frequency distributions also differed between cases and controls (*p* = 0.0028). The proportion of CHD is respectively 46% in CC genotype, 61% in CT genotype and 59% in TT genotype. Frequency of CHD is significantly lower in CC genotype than the other two genotypes, prompting that the loci may associated with coronary heart disease, CC genotype has protective effect on the onset of coronary heart disease. While for other SNP loci, the genotype frequency distributions have no significant difference (Table 4).

**Table 3.**
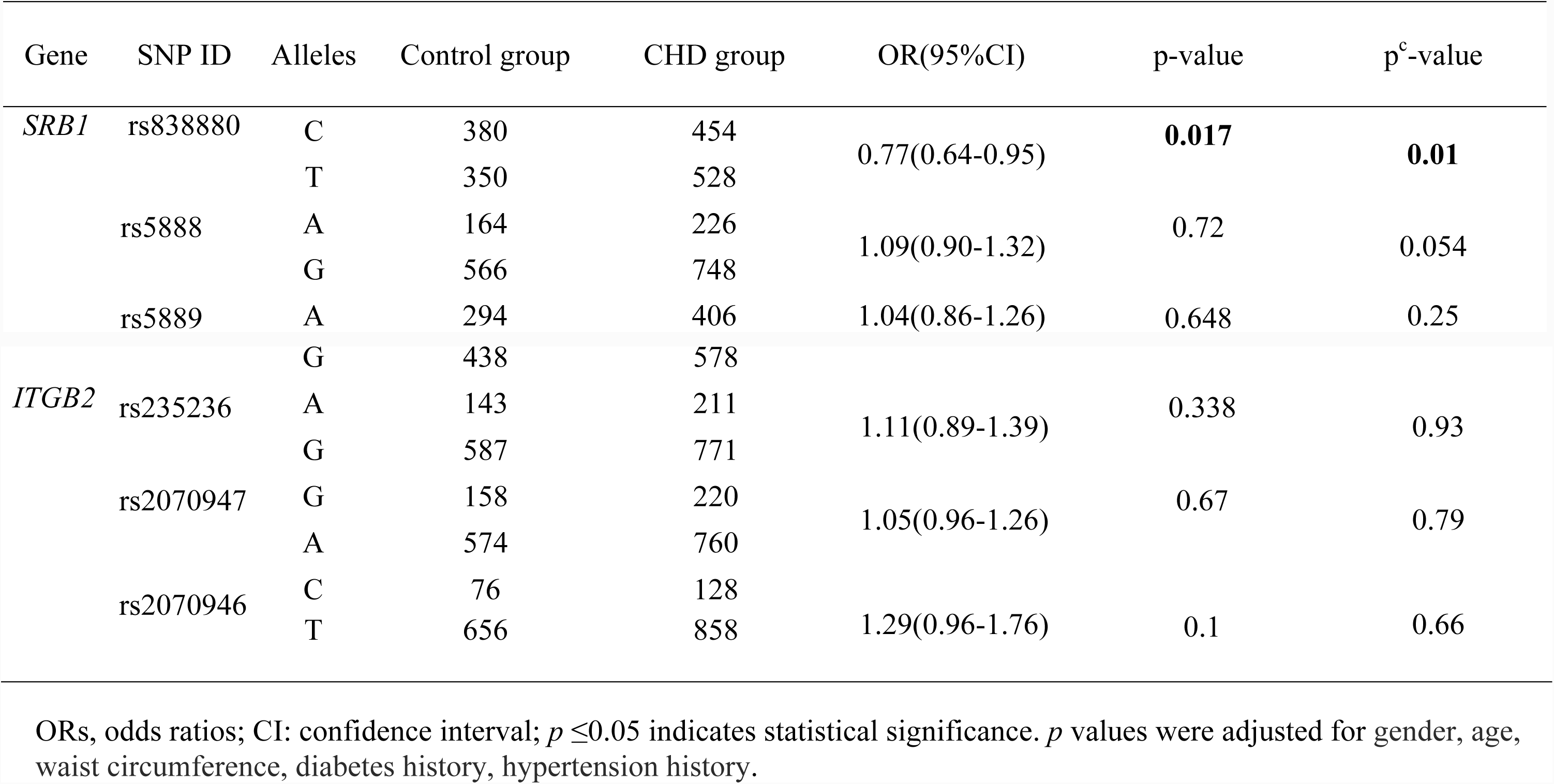
Allele frequencies in cases and controls

**Table 4.**
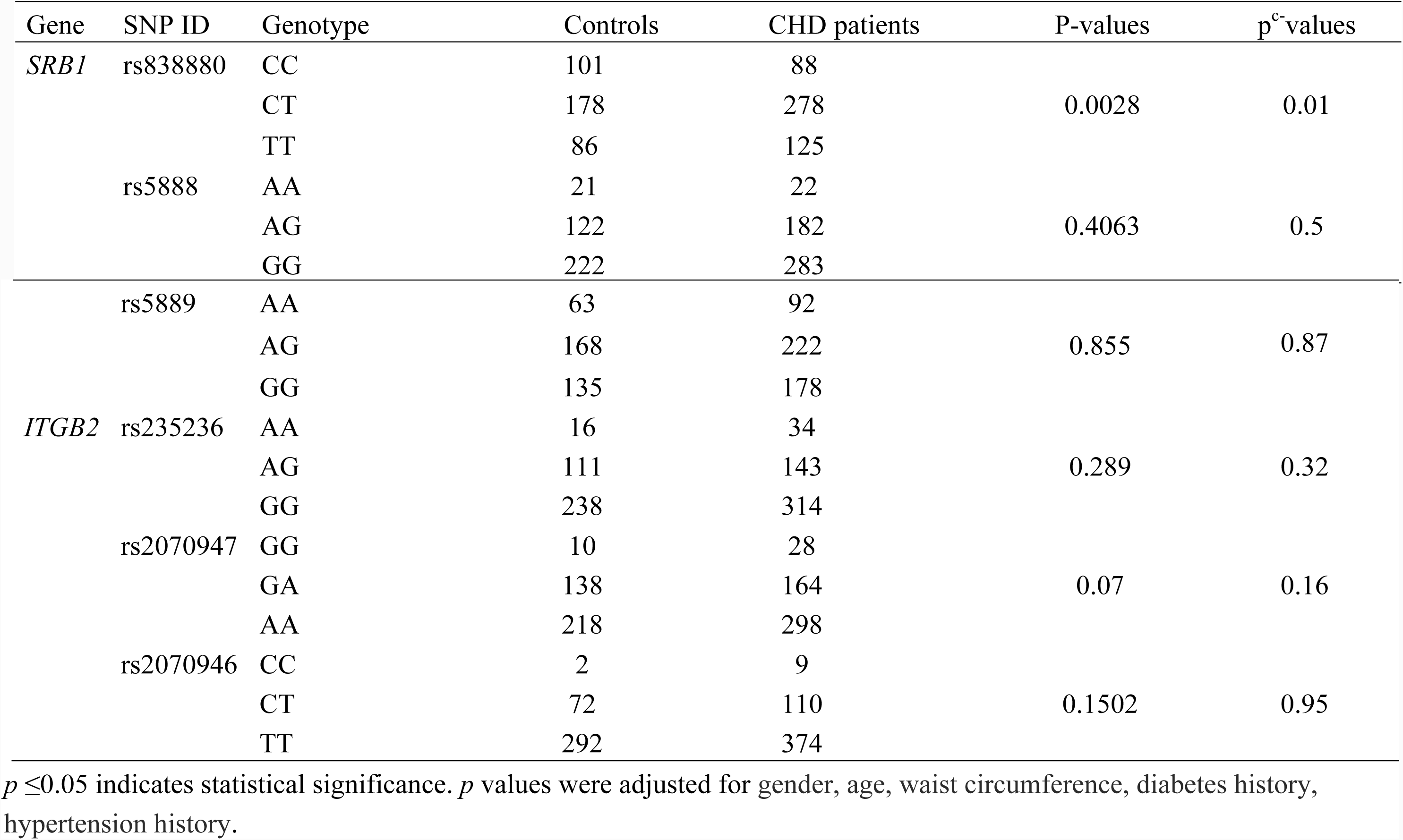
Genotype distribution for SNPs in CHD patients and controls

We assumed the minor allele of each SNP was a dominant or recessive gene loci and analyzed associations between SNPs and CHD in various inheritance models (Table 5–6). According to the results of recessive model, gene SRB1 SNP rs838880 (CC vs CT + TT) have significant difference (p = 0.000672) between two groups, incidence of coronary heart disease of CC genotype group (negative homozygous) was 46%, and CT or TT genotype groups (heterozygous or dominant homozygous) was 62%. Compared with CT or TT type, the CC type has lower incidence of coronary heart disease. Thus rs838880 was associated with a decreased CHD risk in the recessive model. The other two SNP sites of gene SRB1, rs5888 and rs5889, has no statistically significant difference between CHD group and control group (P= 0.4148 and 0.5758 respectively). According to the results of recessive model, gene ITGB2 SNP rs2070947 (GG vs GA + AA) have significant difference (p = 0.03) between two groups, incidence of coronary heart disease of GG genotype group was 73%, and GA or AA genotype groups were 56%. Compared with GA or AA genotype, the GG type has higher incidence of coronary heart disease. Thus rs2070947 was associated with a increased CHD risk in the recessive model. The other SNP sites of gene ITGB2 had no statistically significant difference between CHD group and control group.

**Table 5.** Case-control analysis of dominant model

| Gene | SNP ID | Dominant model | Controls | CHD patients | P-values | P<sup>c</sup>-values |
| --- | --- | --- | --- | --- | --- | --- |
| *SRB1* | rs838880 | CC+CT | 279 | 366 | 0.524 | 0.59 |
|  |  | TT | 86 | 125 |  |  |
|  | rs5888 | AA+AG | 143 | 204 | 0.426 | 0.51 |
|  |  | GG | 222 | 283 |  |  |
|  | rs5889 | AA+AG | 231 | 314 | 0.832 | 0.88 |
|  |  | GG | 135 | 178 |  |  |
| *ITGB2* | rs235236 | AA+AG | 127 | 177 | 0.705 | 0.91 |
|  |  | GG | 238 | 314 |  |  |
|  | rs2070947 | GG+AG | 148 | 192 | 0.711 | 0.75 |
|  |  | AA | 218 | 298 |  |  |
|  | rs2070946 | CC+CT | 74 | 119 | 0.174 | 0,23 |
|  |  | TT | 292 | 374 |  |  |

$p \leq 0.05$ indicates statistical significance. $p$ values were adjusted for gender, age, waist circumference, diabetes history, hypertension history.

**Table 6.** Case-control analysis of recessive model

| Gene | SNP ID | recessive model | Controls | CHD patients | P-value | p<sup>c</sup>-value |
| --- | --- | --- | --- | --- | --- | --- |
| *SRB1* | rs838880 | CC | 101 | 88 | 0.000672 | 0.001 |
|  |  | CT+TT | 246 | 403 |  |  |
|  | rs5888 | AA | 21 | 22 | 0.4148 | 0.5 |
|  |  | AG+GG | 344 | 465 |  |  |
|  | rs5889 | AA | 63 | 92 | 0.5758 | 0.59 |
|  |  | AG+GG | 303 | 400 |  |  |
| *ITGB2* | rs235236 | AA | 16 | 34 | 0.1169 | 0.20 |
|  |  | AG+GG | 349 | 457 |  |  |
|  | rs2070947 | GG | 10 | 28 | 0.03611 | 0.09 |
|  |  | GA+AA | 356 | 462 |  |  |
|  | rs2070946 | CC | 2 | 9 | 0.09918 | 0.15 |
|  |  | CT+TT | 364 | 484 |  |  |

$p \leq 0.05$ indicates statistical significance. $p$ values were adjusted for gender, age, waist circumference, diabetes history, hypertension history.

We compared the clinical and serum metabolic indexes of SNP sites by genotype (Table 7). We found significant differences in the history of dyslipidemia, LPA among different genotypes of SRB1 rs5888 (*p* < 0.05). Subjects carrying the rs5888 AA genotype had lower history of hyperlipidemia and LPA level than those carrying the AG and GG genotypes. We observed a significant difference in hypertension, HDL, APOA1 among gene ITGB2 rs235236 genotypes (*p* < 0.05). Patients who carried the AA genotype displayed higher HDL levels and patients who carried the AG genotype displayed lower APOA1 levels. In gene ITGB2 rs2070947, TG level is higher in GA genotype (p=0.01). We also observed a significant difference in levels of APOA1 and hsCRP among the ITGB2 rs2070946 genotypes (*p* < 0.05). Patients who carried the CT genotype displayed higher APOA1 and hsCRP levels.

**Table 7.**
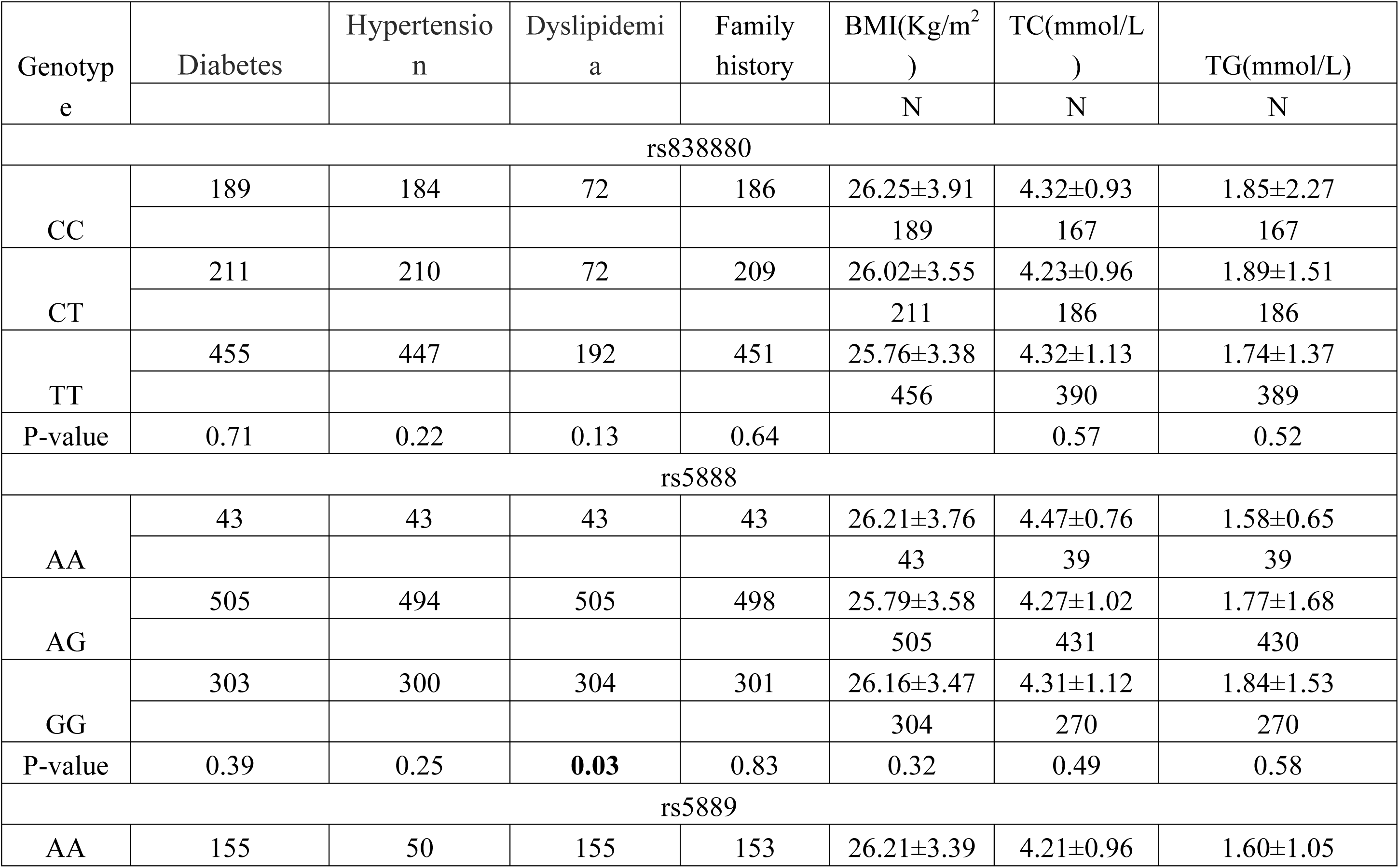
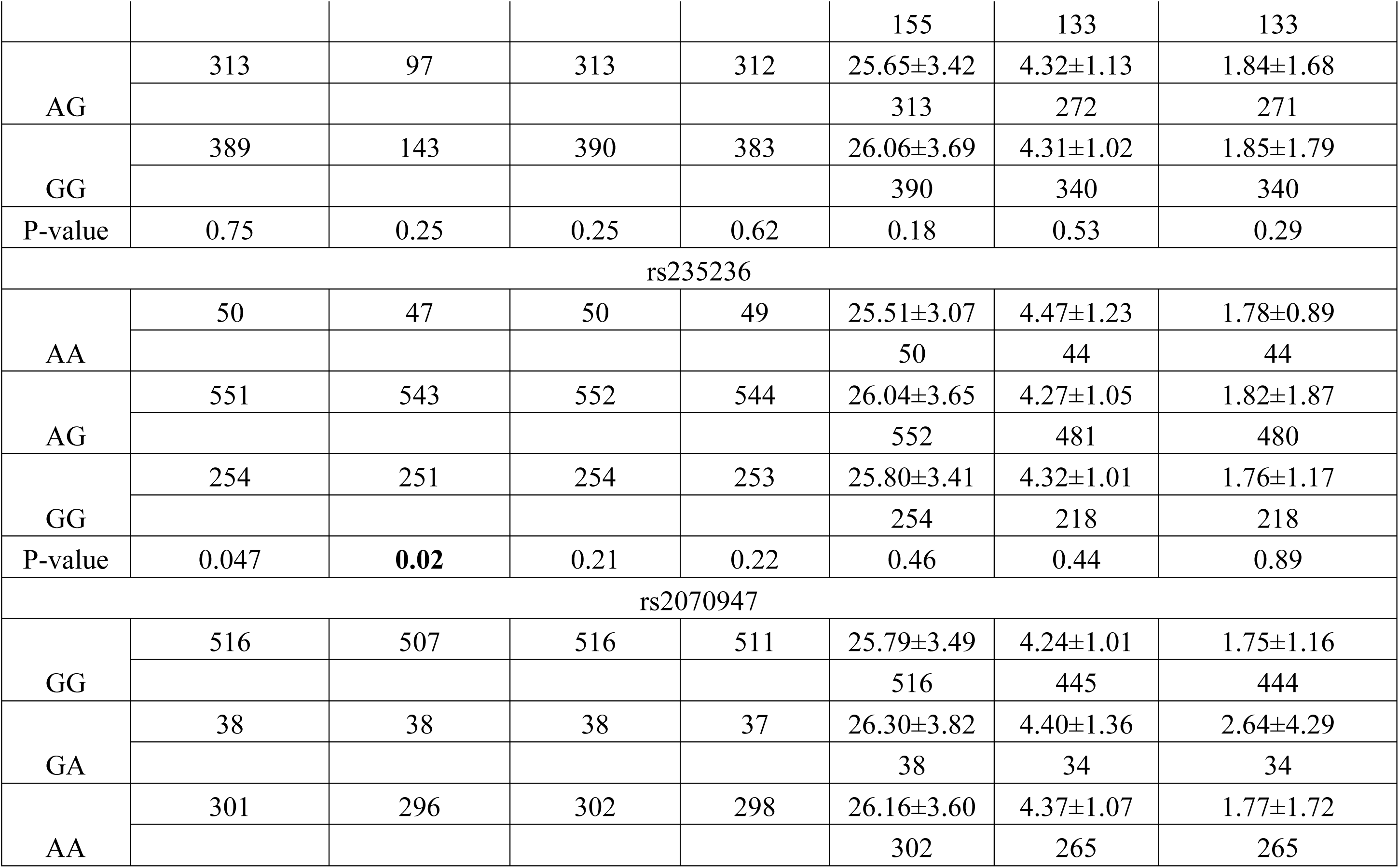
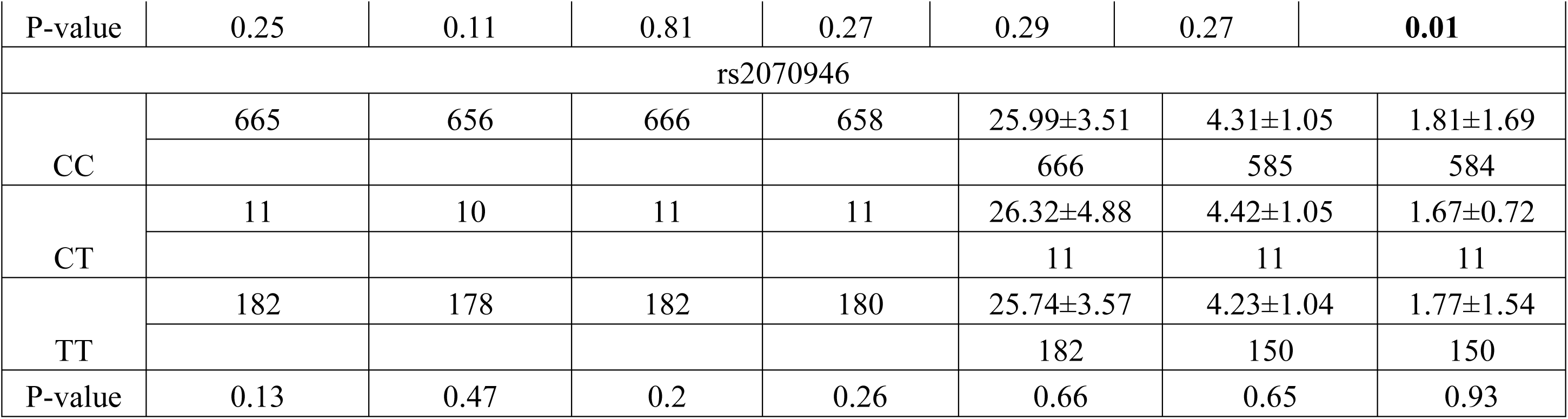
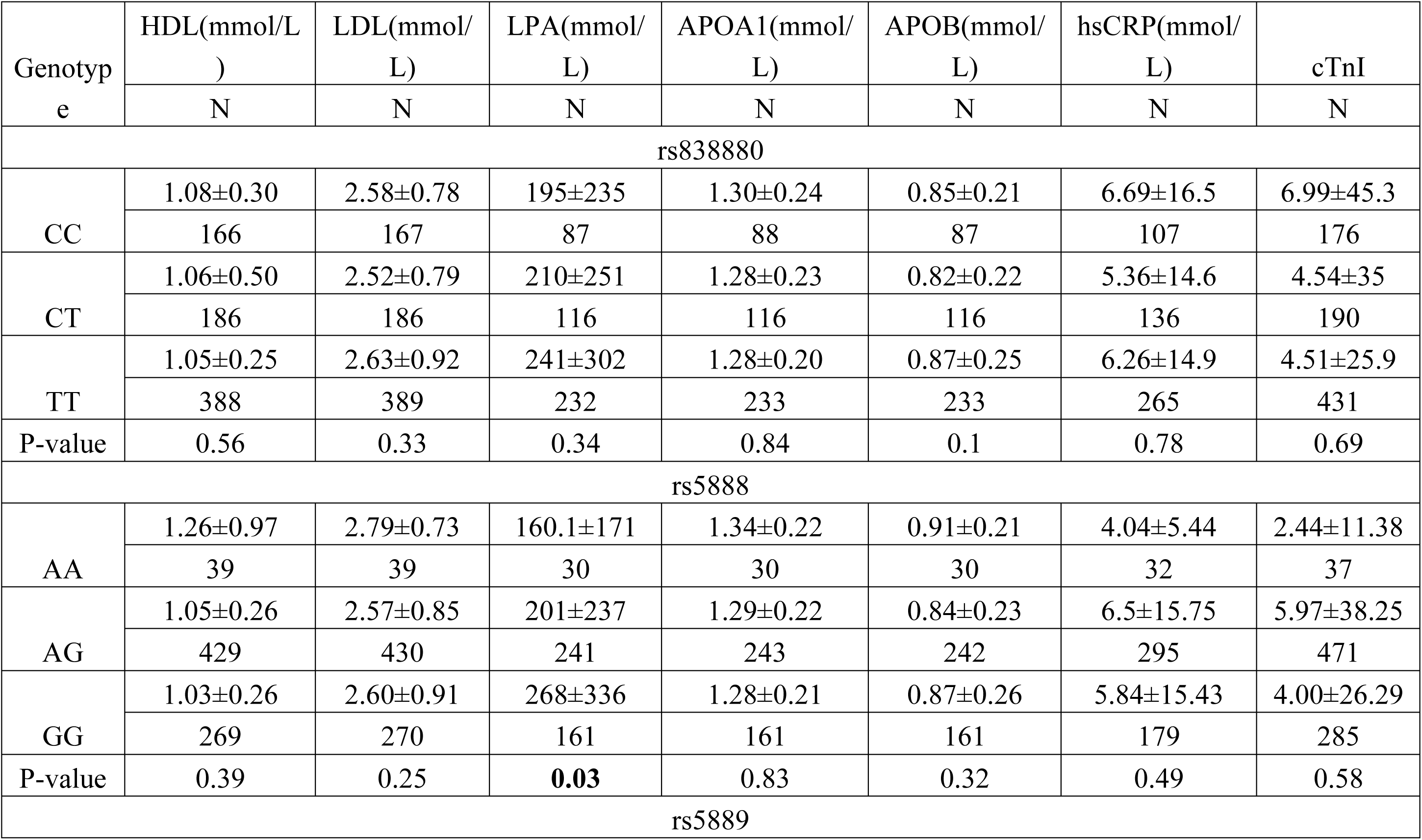
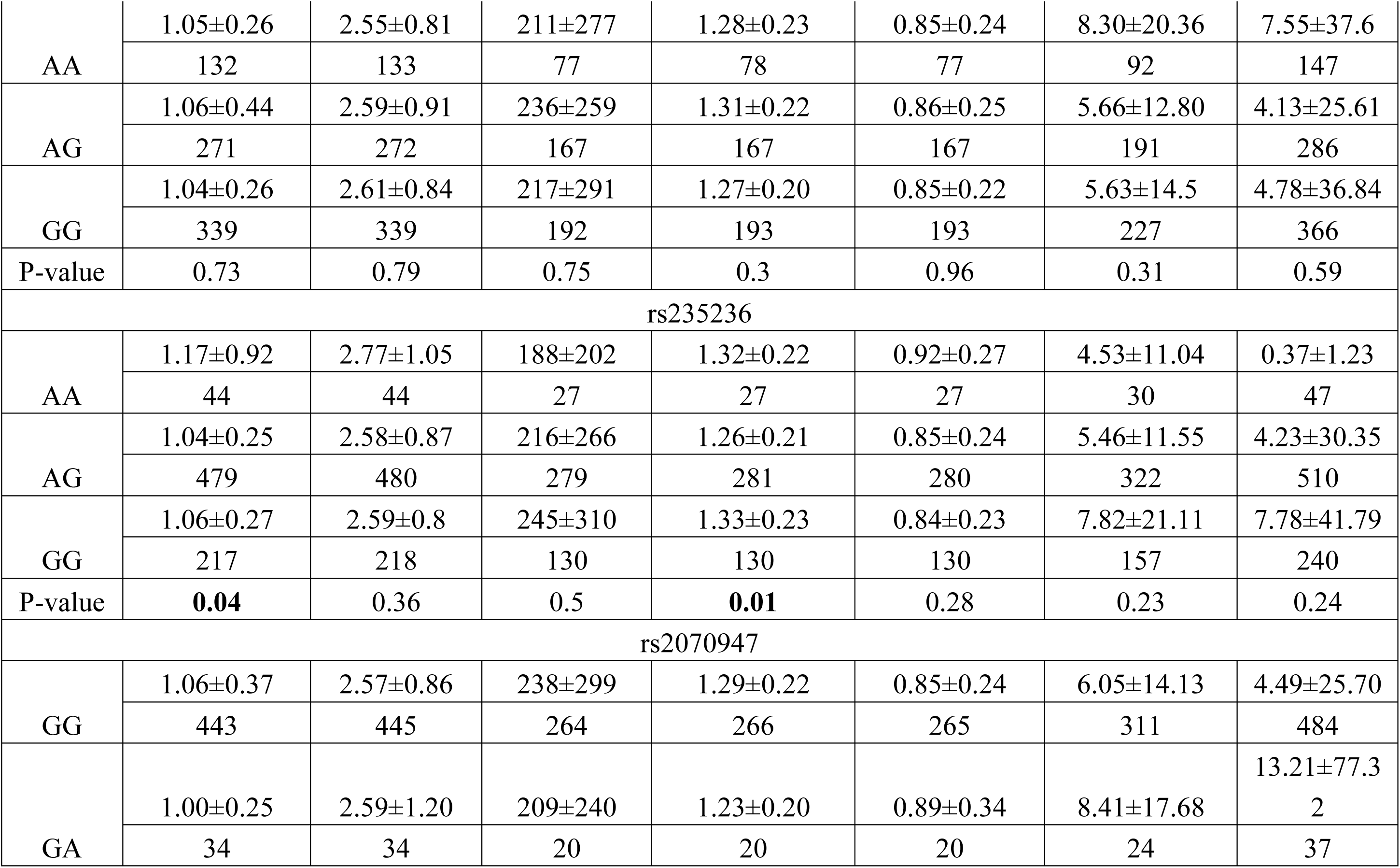
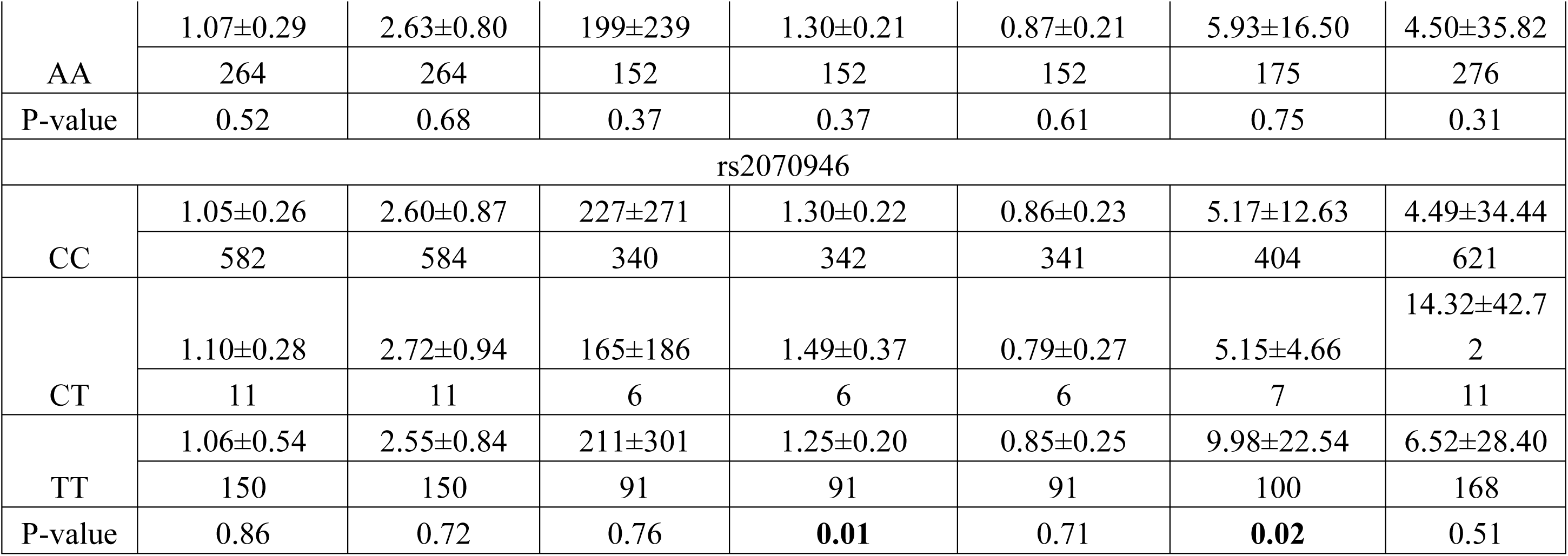
Clinical and biochemical characteristics of participants

**Table 8.** *ITGB2* haplotype frequencies and the association with the risk of CHD

| Block | Frequency | Case, Control Ratio | P Value |
| --- | --- | --- | --- |
| AT | 66.1% | 0.646, 0.681 | 0.1276 |
| GT | 22% | 0.224, 0.215 | 0.647 |
| AC | 11.9% | 0.130, 0.104 | 0.1006 |

$p \leq 0.05$ indicates statistical significance.

Haplotype analysis revealed one block in ITGB2 (Figure 1): including rs2070947 and rs2070946. Further analyses of associations between ITGB2 haplotypes and CHD risk had a negative result.

**Figure 1:**
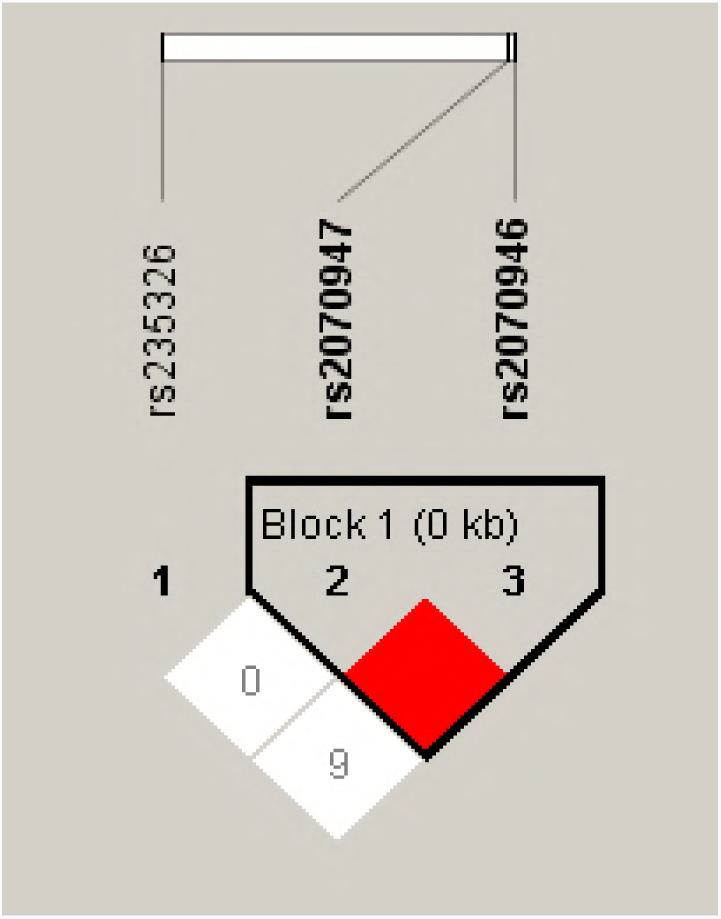
Haplotype block map for all the SNPs of the *ITGB2* gene.

## Discussion

Coronary heart disease (CHD) is a complex disease caused by long-term interaction between several micro-acting genes and environmental factors. In recent years, a series of genes or mutations associated with coronary heart disease have been found worldwide through the whole genome association study. Development of coronary heart disease (CHD) genome research to parse the genetic and molecular mechanism of CHD and myocardial infarction has important scientific value for early warning of CHD, early screening and forecast of high-risk population, clinical diagnosis and new drug research and development.

The selecting of control group and case group is critical for genomic analysis. Because any systematic error between two groups can result in false-positive, false-negative results. There are many risk factors for coronary heart disease. Mainly includes the following aspects: age, with the increase of age, the incidence of coronary heart disease increase; gender, male has higher cardiovascular disease rates than female; diabetes, the risk of cardiovascular disease is significant increase in patients with diabetes or glucose intolerance; dyslipidemia, elevated serum total cholesterol and low density lipoprotein are risk factors for coronary heart disease and ischemic stroke. Other risk factors include smoking, high blood pressure, and family history of cardiovascular disease. The research subjects were selected from the database established by the CHD research group from 2009. All the subjects were examined by coronary angiography in our hospital and obtained detailed clinical data. To avoid systematic errors caused by inconsistent of general situation and basic diseases between the two groups, coronary heart disease group and control group were matched in age, gender, BMI, diabetes history, dyslipidemia. Because of the limitations of sample size, smoking and drinking history and history of hypertension were not matched. Coronary angiography is the gold standard in the diagnosis of coronary heart disease, and it is also our standard of classification. Drawback is that most people undergone coronary angiography has symptoms like chest tightness or chest pain, so our control group is not completely normal crowd. whether this has any impact to our study is not known.

Coronary heart disease (CHD) is a kind of atherosclerotic disease, the occurrence of atherosclerosis (As) has close relationship with lipid metabolic abnormalities. Lower serum high-density lipoprotein (HDL) is an important risk factor of cardiovascular disease ^[22]^. HDL plays a role in anti-atherosclerosis by promoting cholesterol in the arterial wall cells transferred to the liver. Scavenger receptor B type 1 (SRB1) is the primary HDL receptor and a critical receptor in HDL metabolism. SRB1 is widely distributed in liver and tissues secrete steroid hormones, and is also expressed in arterial wall cells, including endothelial cells, smooth muscle cells and macrophages ^[24]^. SRB1 can help maintain the balance of cholesterol metabolism and prevent free cholesterol accumulation in the arterial wall by promoting free cholesterol ransferred from peripheral tissue cell to HDL, mediating the selective uptake of cholesterol in high-density lipoprotein of liver and promoting the secretion of cholesterol in the liver cells into the bile. The absence of SRB1 can result in unbalance of cholesterol, accumulation of lipids in the arterial wall and eventually lead to atherosclerosis ^[16]^. In the experiments of Rigotti et al, the serum total cholesterol levels of the mice with SRB1 gene knockout were significantly higher than those of normal mice and severe atherosclerosis appeared ^[25–27]^. Therefore, SRB1 gene may play a protective role in cardiovascular disease by affecting lipid metabolism. The correlation between single nucleotide polymorphisms of SRB1 gene and dyslipidemia, atherosclerosis and coronary heart disease has been reported. Ani Manichaikul et al. found that rs10846744 of gene SRB1 was significantly associated with atherosclerosis and acute myocardial infarction ^[28]^. In our study, gene SRB1 rs838880 locus allele frequency, genotype frequency and recessive models between the two groups were statistically significant. Indicate that the loci is associated with coronary heart disease (CHD) and C allele have protective effect on coronary heart disease (CHD). Gene SRB1 rs838880 site has been reported related to HDL in foreign reports ^[30]^. There is no research on the correlation between rs838880 site and coronary heart disease up to now. SRB1 genes affect the occurrence and development of coronary heart disease mainly through affects lipid metabolism process. But in our study, there is no statistical differences between three genotypes of SRB1 rs838880 gene loci and clinical and serum metabolism index, including lipid metabolism related indicators. This may be because a large portion of patients with coronary heart disease (CHD) are taking statins (56%), which may have a negative effect on lipid metabolism, and leading to false negative results. The correlation of gene SRB1 rs5888 loci and coronary heart disease has been reported. In study of Daiva, gene SRB1 rs5888 loci was associated with lipid metabolism and coronary heart disease, and associated with the acute myocardial infarction of elderly men ^[29]^. In our study, rs5888 loci has no correlation with coronary heart disease, but is associated with abnormal lipid metabolism. AA genotype has a lower frequency of hyperlipidemia history than the other two genotypes (GG,AG). At the same time, LPA level in AA genotype is significantly lower than the other two genotypes.

It is suggested that the A allele may be a protective factor for the lipid metabolism. In our study, Gene SRB1 rs5889 loci has no relationship with coronary heart disease, and this is consistent with Jeffrey’s results ^[31]^.

Inflammation is one important mechanism of the onset of atherosclerosis. Some risk factors like smoking, high blood glucose promote the onset of the AS by stimulating vascular inflammation. Inflammatory cells composition in the fibrous cap and release of cytokines, inflammatory factors and ultimately lead to plaque rupture, causing acute coronary syndrome (ACS) ^[32]^. The basic of the interaction between leukocyte and endothelial cells is cell adhesion and adhesion molecule ^[17]^. Therefore adhesion molecules play an important role in the inflammatory process. Integrin beta 2 (ITGB2), namely CD18, is an important member of the adhesion molecule integrin family. ITGB2 expression mainly on neutrophils and monocytes. It mediates the interaction between leukocyte and endothelial cells through adhesion molecules on endotheliocyte like intercellular adhesion molecule-1(ICAM1). In our early proteomics research (Study on proteomics of epicardial fat and subcutaneous fat in coronary heart disease patients). It has been confirmed that the expression of ITGB2 is different in CHD and non-CHD (both in the epicardial fat and subcutaneous fat). Expression level of ITGB2 rise in CHD group (1.6 times of non-CHD group) and reduce in subcutaneous fat (0.1 times of non-CHD group) ^[18]^. Epicardial fat has close relationship with coronary heart disease (CHD). The inflammatory response in epicardial fat is significantly stronger than in subcutaneous fat in severe coronary heart disease patients and we can find different sources of inflammatory cells infiltration such as lymphocytes, macrophages, mast cells through immunohistochemical ^[33]^. group. The expression of ITGB2 increased in epicardial fat in CHD. Indicating that ITGB2 may participate in the cell adhesion and chemotaxis thus palys a role in the inflammatory process of coronary artery. Michael Kassirer examined CD11b/CD18 antigen expression on peripheral blood leukocyte in patients with ischemic heart disease, healthy controls and posttraumatic patients (no cardiovascular disease). Result is that CD11b/CD18 antigen expression significantly increased in ischemic heart disease patients. Indicating that there is an activation of peripheral blood leukocyte in ischemic heart disease patients and the activation is mainly reflected the atherosclerotic plaque inflammation rather than caused by acute stress ^[34]^. The evidence suggests that ITGB2 affects the occurrence and development of coronary heart disease through inflammatory response mechanisms. But there is less research on the relationship between single nucleotide polymorphisms and coronary heart disease in the gene ITGB2. The study combined our earlier proteomics results, which is a innovation of this study. However, our positive result is not much in correlation analysis. This maybe because the SNP site we select is relatively new. There is less study about ITGB2 SNP loci and cardiovascular disease. In addition, there are certain limitations of correlation studies due to the stratification, regional differences and heterogeneity of genetic background. To investigate whether genetic polymorphism is associated with coronary heart disease, it’s need to expand the sample size and to verify in different populations.

## Conclusion

Gene SRB1 rs838880 loci (C/T) has correlation with the occurrence of coronary heart disease, CC genotype has lower risk of coronary heart disease, C allele might be protective in the onset of coronary heart disease (CHD). Gene SRB1 rs5888 has relationship with lipid metabolism, and A allele may be protective in lipid metabolism. Gene ITGB2 rs2070947 loci, GG genotype has higher incidence of coronary heart disease (73%) than that of genotype for GA or AA type (56%), GG genotype may be a risk factor for coronary heart disease. Gene ITGB2 rs2070946 CT genotype has elevated hsCRP level, which may be a risk factor for the inflammatory response of coronary heart disease.

